## Supplemental Legends for "Avian-specific *Salmonella* transition to endemicity is accompanied by localized resistome and mobilome interaction"

### Supplementary figures:

**S1 Fig. The evolutionary structure of global *S*. Gallinarum.** The phylogenetic tree was constructed using cgSNPs, revealing three *S.* Gallinarum biovars: bvSP (n=528/580, 91.03%), bvSG (n=50/580, 8.6%), and bvSD (n=2/580, 0.34%). Additionally, *Salmonella* serovar Enteritidis (SE) is represented by a gray line. Employing hierarchical Bayesian analysis, bvSP was further subdivided into five lineages: fuchsia (L1), orange (L2a), pink (L2b), red (L3b), and green (L3c). The colorful circles indicate detailed information on *Salmonella* sequence type and isolation regions. The outermost circle denotes the locations of the 45 bvSP strains isolated from Yueqing and Taishun.

**S2 Fig. The primary prevalence of bvSP lineages varies across different regions of China over time, with consideration given to the eastern, northern, and southern regions.** The colors indicate the specific lineage type prevalence within each region: fuchsia (L1), orange (L2a), pink (L2b), red (L3b), and green (L3c).

**S6 Fig. Assessment of the Temporal Structure (L1-L3c).** The plots depict the root-to-tip regression analysis for the *Salmonella* maximum likelihood tree, generated using Treetime software. Each data point on the plot represents a measurement from the root to each tip in the tree, with the solid line indicating the regression line.

**S7 Fig. Historical international transmissions of bvSP lineages L2b and L3b are depicted with arrows representing the transmission paths. The pink and red lines represent L2b and L3b, respectively.**

**S8 Fig. Potential transmission events (n=53) of *S*. Gallinarum biovar Pullorum (bvSP).** The cgSNP distances were calculated between the bvSP strains isolated from Zhejiang Province (n=95) and those from China with available provincial information (n=435). Only cgSNP distances less than two are depicted, with darker colors indicating a higher transmission event.

**S9 Fig. The carriage of four predominant mobilome.** (a). The phylogenetic tree of *S*. Gallinarum was constructed using cgSNPs, with distinct colors representing each *S*. Gallinarum biovar; *Salmonella serovar* Enteritidis (SE.) is depicted in gray. Furthermore, different colors are assigned to represent various lineages of bvSP: fuchsia for L1, orange for L2a, pink for L2b, red for

L3b, and green for L3c. Heatmaps on the right side illustrate the presence of integrons, transposons, plasmids, and prophages carried by the corresponding *Salmonella* strains. (b). Predominant types of mobilomes prevalent among bvSP are depicted. The x-axis of the bar graph illustrates the top five mobilome types in bvSP based on the total count within each category.

**S10 Fig. Types of predominant mobile genetic elements carried by various regions.** From a-d presents integron, transposon, plasmid, and prophage, respectively.

**S11 Fig. Trends in both resistome and mobilome quantities over time and across lineages.**

**S12 Fig. Recombination removal using Gubbins.** Recombination in five lineages (L1, L2a, L2b, L3b, L3c) were removed using Gubbins with default parameters. The recombination regions for each lineage were mapped onto the reference genome, *S*. Gallinarum R51. Different colors represent the number of recombination events in each *S*. Gallinarum lineages strains, with darker colors indicating higher frequencies of recombination.

### Supplementary tables:

**S1 Table.** Information on 45 newly isolated bvSP originated from Yueqing and Taishun used in this study.

**S2 Table.** Information on 540 *Salmonella* isolates was obtained from public sources to assemble the global database, with 325 sequences previously preserved in our laboratory.

**S3 Table.** The regional classification of 436 bvSP strains isolated from China was conducted.

**S4 Table.** Information on calculation of invasiveness index for 45 bvSP isolates newly originated from Yueqing and Taishun

**S5 Table.** SNP distance-based tracing analysis for the 95 strains from Zhejiang Province and those from China with available provincial information (n=435). Only strains with an SNP distance of two or fewer are considered likely to be involved in potential transmission events.

**S6 Table.** Information on antimicrobial resistance genes carried by 528 bvSP isolates.

**S7 Table.** Information on plasmids, transposons, integrons, and prophages carried by 528 bvSP isolates.

**S8 Table.** A co-localization analysis was conducted to assess each ARG's association with mobile genetic elements (MGEs). Among 621 ARGs identified in 436 bvSP isolates collected across China, 415 ARGs were found to be located on MGEs.

**S9 Table.** Detection of Horizontal Gene Transfer (HGT) of ARGs Carried by mobile genetic elements in bvSP Genomes from China. Using the HGTphyloDetect pipeline, we calculated the Alien Index (AI) score and out_perc values for each ARG sequences. ARGs with AI score ≥ 45 and out_perc ≥ 90% were identified as potential candidates for horizontal ARGs transfer. Additionally, based on BLAST hit scores, we determined the most likely donor organisms for these ARGs.

**S10 Table.** The HGT frequency value for specific antimicrobial resistance genes was identified from bvSP isolated from different regions of China.
