## Supplemental figures 1-12 for "Avian-specific *Salmonella* transition to endemicity is accompanied by localized resistome and mobilome interaction"

### Supplementary figures

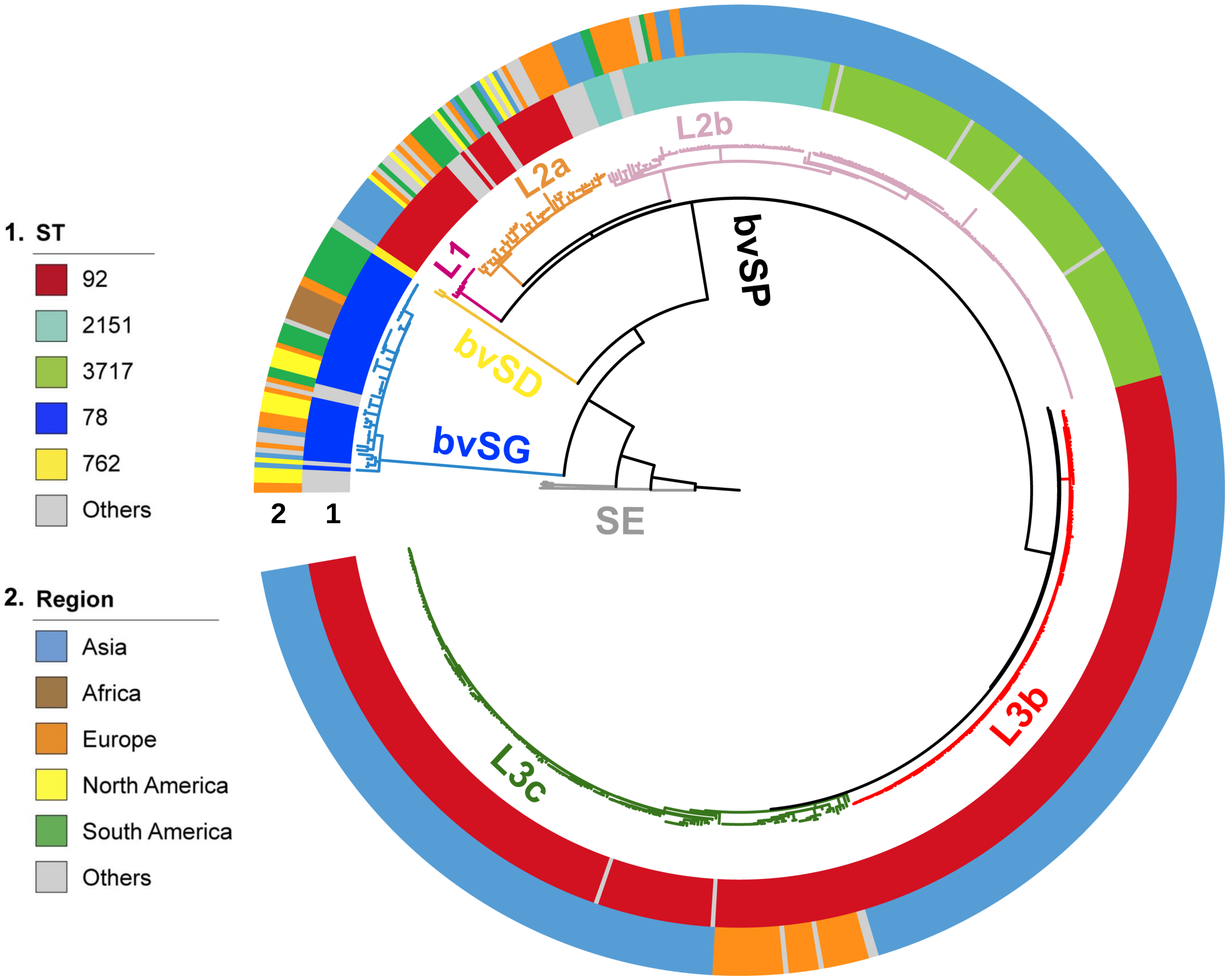


**S1 Fig. The evolutionary structure of global *S*. Gallinarum.** The phylogenetic tree was constructed using cgSNPs, revealing three *S.* Gallinarum biovars: bvSP (n=528/580, 91.03%), bvSG (n=50/580, 8.6%), and bvSD (n=2/580, 0.34%). Additionally, *Salmonella* serovar Enteritidis (SE) is represented by a gray line. Employing hierarchical Bayesian analysis, bvSP was further subdivided into five lineages: fuchsia (L1), orange (L2a), pink (L2b), red (L3b), and green (L3c). The colorful circles indicate detailed information on *Salmonella* sequence type and isolation regions. The outermost circle denotes the locations of the 45 bvSP strains isolated from Yueqing and Taishun.

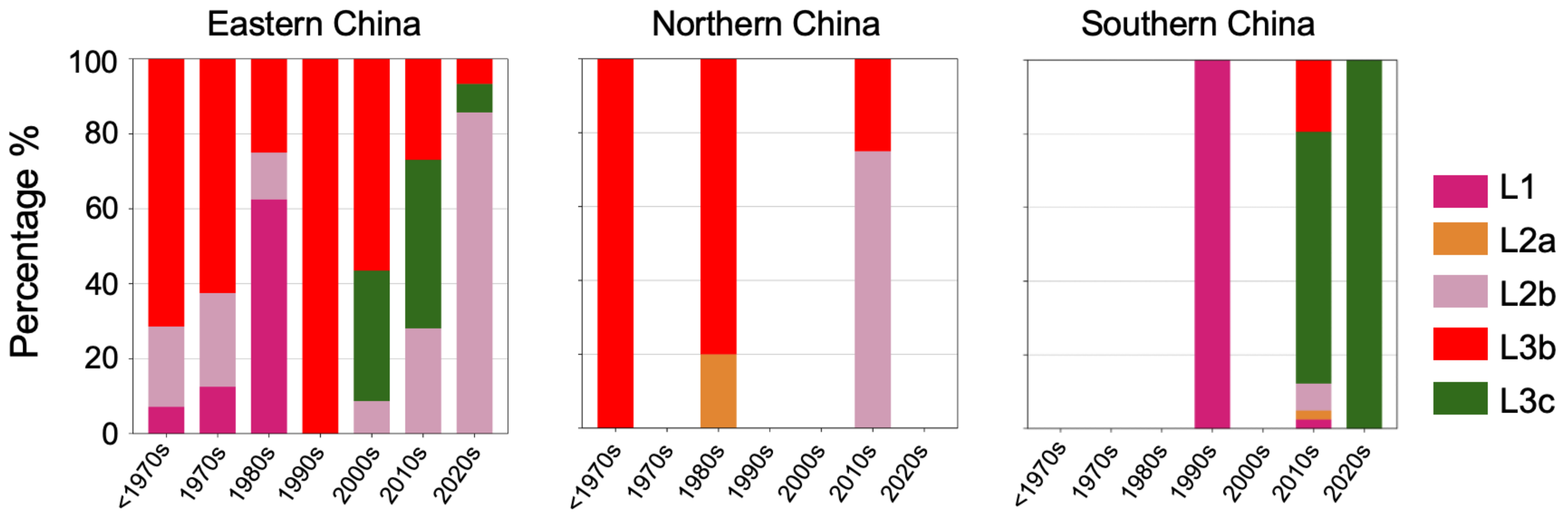


**S2 Fig. The primary prevalence of bvSP lineages varies across different regions of China over time, with consideration given to the eastern, northern, and southern regions.** The colors indicate the specific lineage type prevalence within each region: fuchsia (L1), orange (L2a), pink (L2b), red (L3b), and green (L3c).

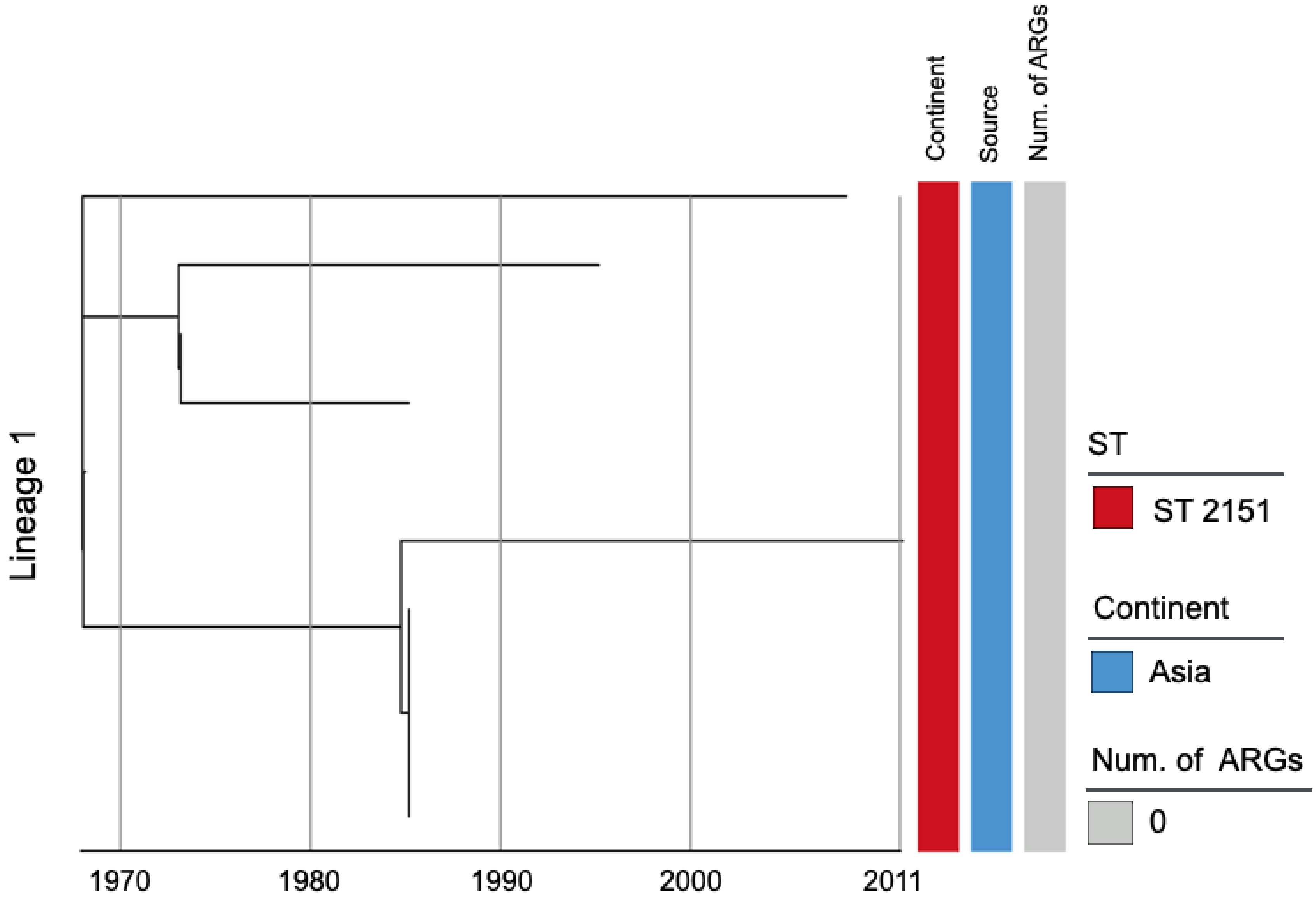


**S3 Fig. Phylogenetic tree of *Salmonella* Gallinarum L1 based on a spatiotemporal Bayesian framework.** The phylogenetic tree on the left was constructed using a reference-mapped multiple core-genome SNPs sequence alignment, with recombination regions detected and removed by Gubbins. The spatiotemporal Bayesian framework was configured with the "GTR" substitution model, 4 Gamma Category Count, "Relaxed Clock Log Normal" model, "Coalescent Bayesian Skyline" tree prior model, and a Markov Chain Monte Carlo (MCMC) chain length of 100,000,000, with sampling every 10,000 iterations. Convergence was assessed using Tracer, ensuring all parameter effective sampling sizes (ESS) exceeded 200. Evolutionary time is represented by the length of the branches. The heatmap on the right displays, respectively, the sequence type (ST), region of isolation, and the number (Num.) of antimicrobial resistance genes (ARGs) carried by the corresponding *Salmonella* Gallinarum.

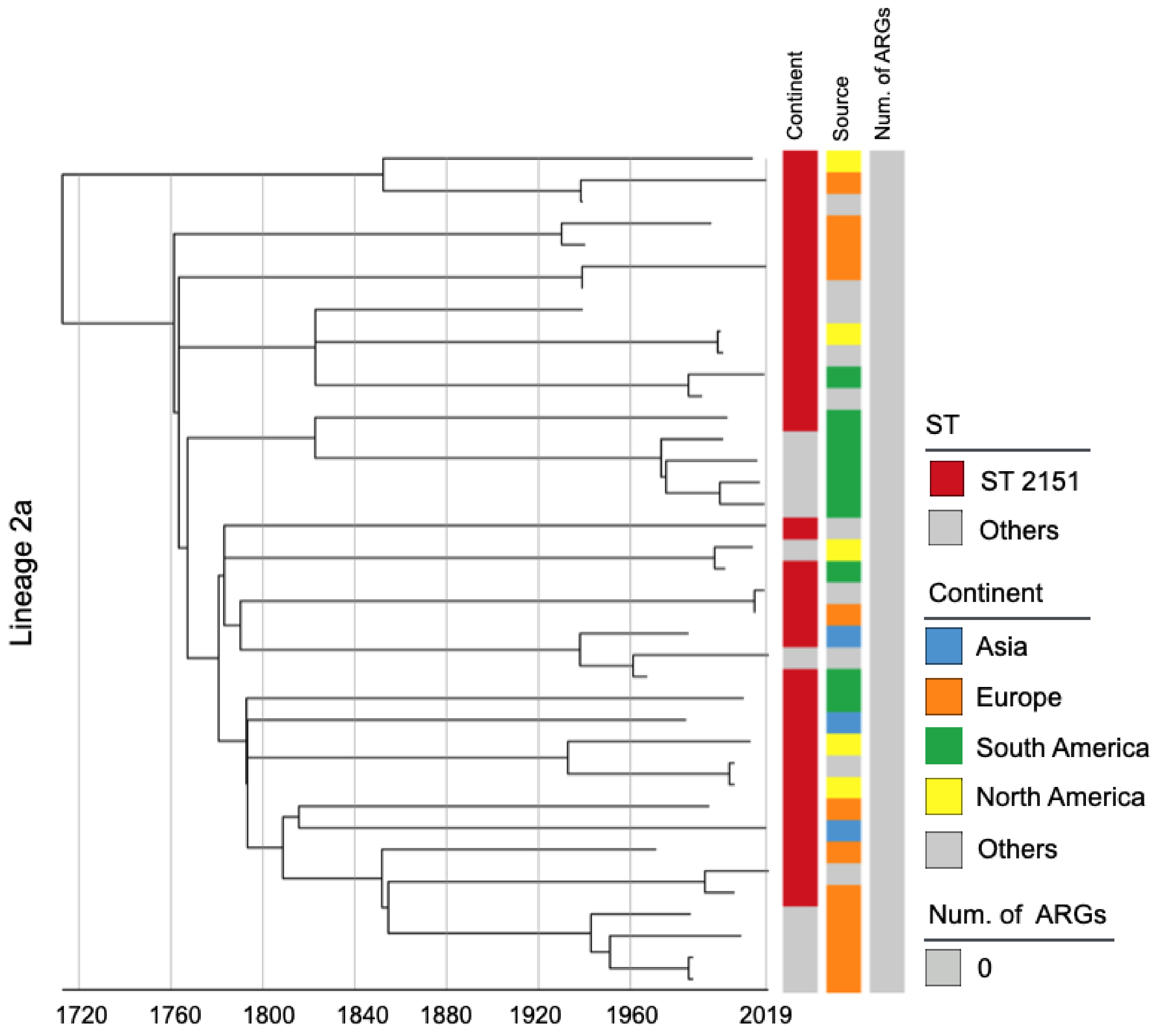


**S4 Fig. Phylogenetic tree of *Salmonella* Gallinarum L2a based on a spatiotemporal Bayesian framework.** The phylogenetic tree on the left was constructed using a reference-mapped multiple core-genome SNPs sequence alignment, with recombination regions detected and removed by Gubbins. The spatiotemporal Bayesian framework was configured with the "GTR" substitution model, 4 Gamma Category Count, "Relaxed Clock Log Normal" model, "Coalescent Bayesian Skyline" tree prior model, and a Markov Chain Monte Carlo (MCMC) chain length of 100,000,000, with sampling every 10,000 iterations. Convergence was assessed using Tracer, ensuring all parameter effective sampling sizes (ESS) exceeded 200. Evolutionary time is represented by the length of the branches. The heatmap on the right displays, respectively, the sequence type (ST), region of isolation, and the number (Num.) of antimicrobial resistance genes (ARGs) carried by the corresponding *Salmonella* Gallinarum.

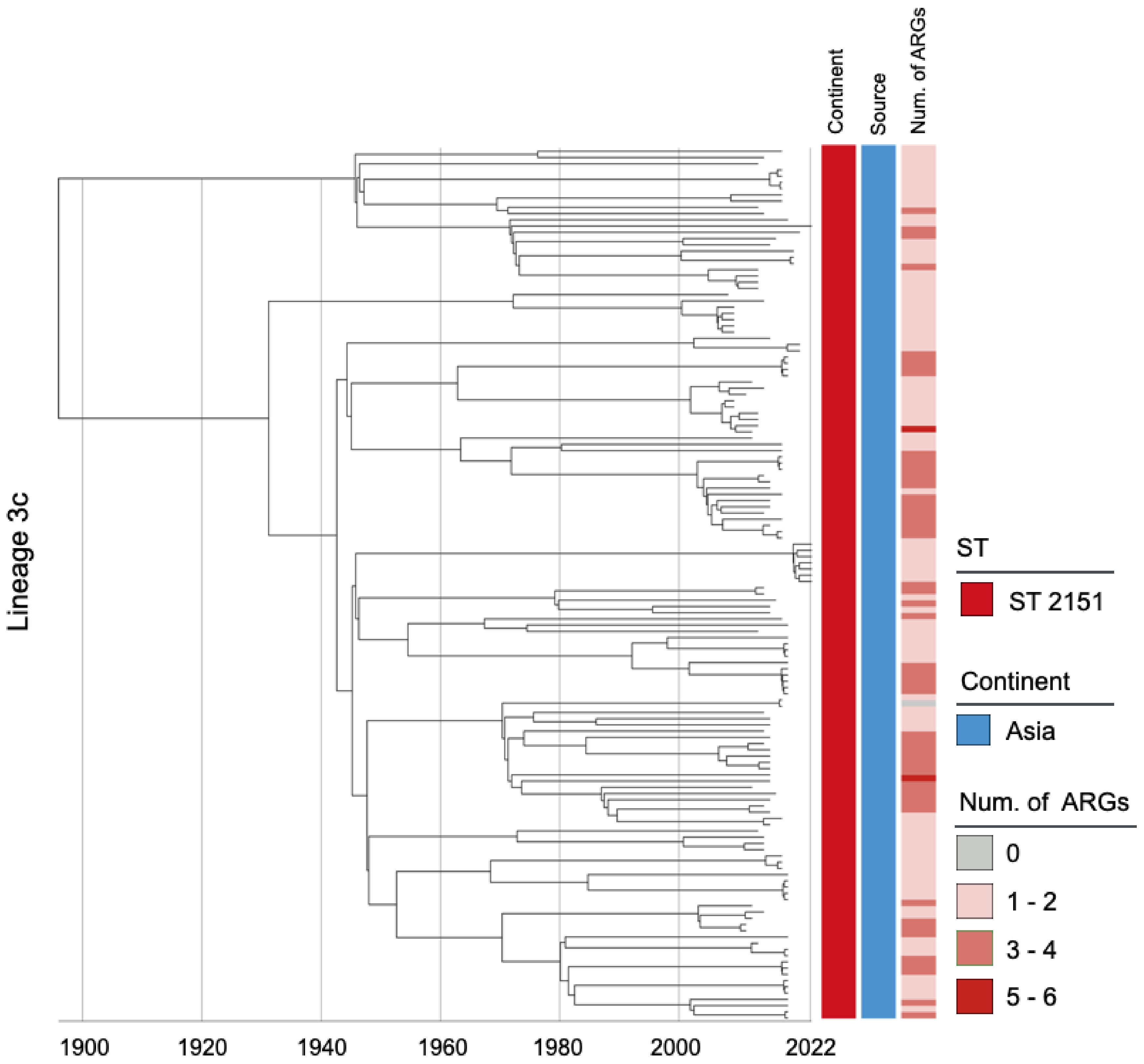


**S5 Fig. Phylogenetic tree of *Salmonella* Gallinarum L3c based on a spatiotemporal Bayesian framework.** The phylogenetic tree on the left was constructed using a reference-mapped multiple core-genome SNPs sequence alignment, with recombination regions detected and removed by Gubbins. The spatiotemporal Bayesian framework was configured with the "GTR" substitution model, 4 Gamma Category Count, "Relaxed Clock Log Normal" model, "Coalescent Bayesian Skyline" tree prior model, and a Markov Chain Monte Carlo (MCMC) chain length of 100,000,000, with sampling every 10,000 iterations. Convergence was assessed using Tracer, ensuring all parameter effective sampling sizes (ESS) exceeded 200. Evolutionary time is represented by the length of the branches. The heatmap on the right displays, respectively, the sequence type (ST), region of isolation, and the number (Num.) of antimicrobial resistance genes (ARGs) carried by the corresponding *Salmonella* Gallinarum.

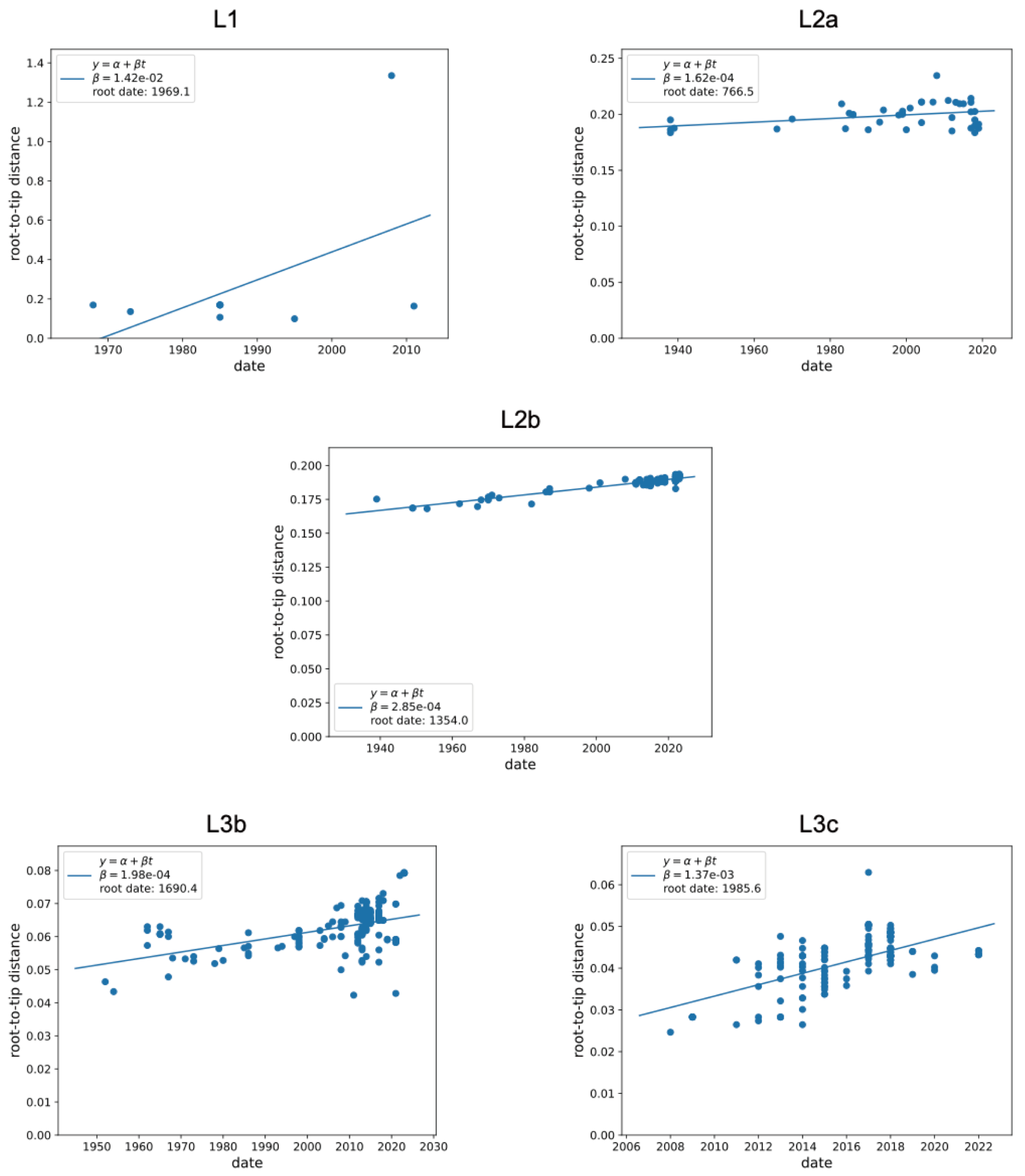


**S6 Fig. Assessment of the Temporal Structure (L1-L3c).** The plots depict the root-to-tip regression analysis for the *Salmonella* maximum likelihood tree, generated using Treetime software. Each data point on the plot represents a measurement from the root to each tip in the tree, with the solid line indicating the regression line.

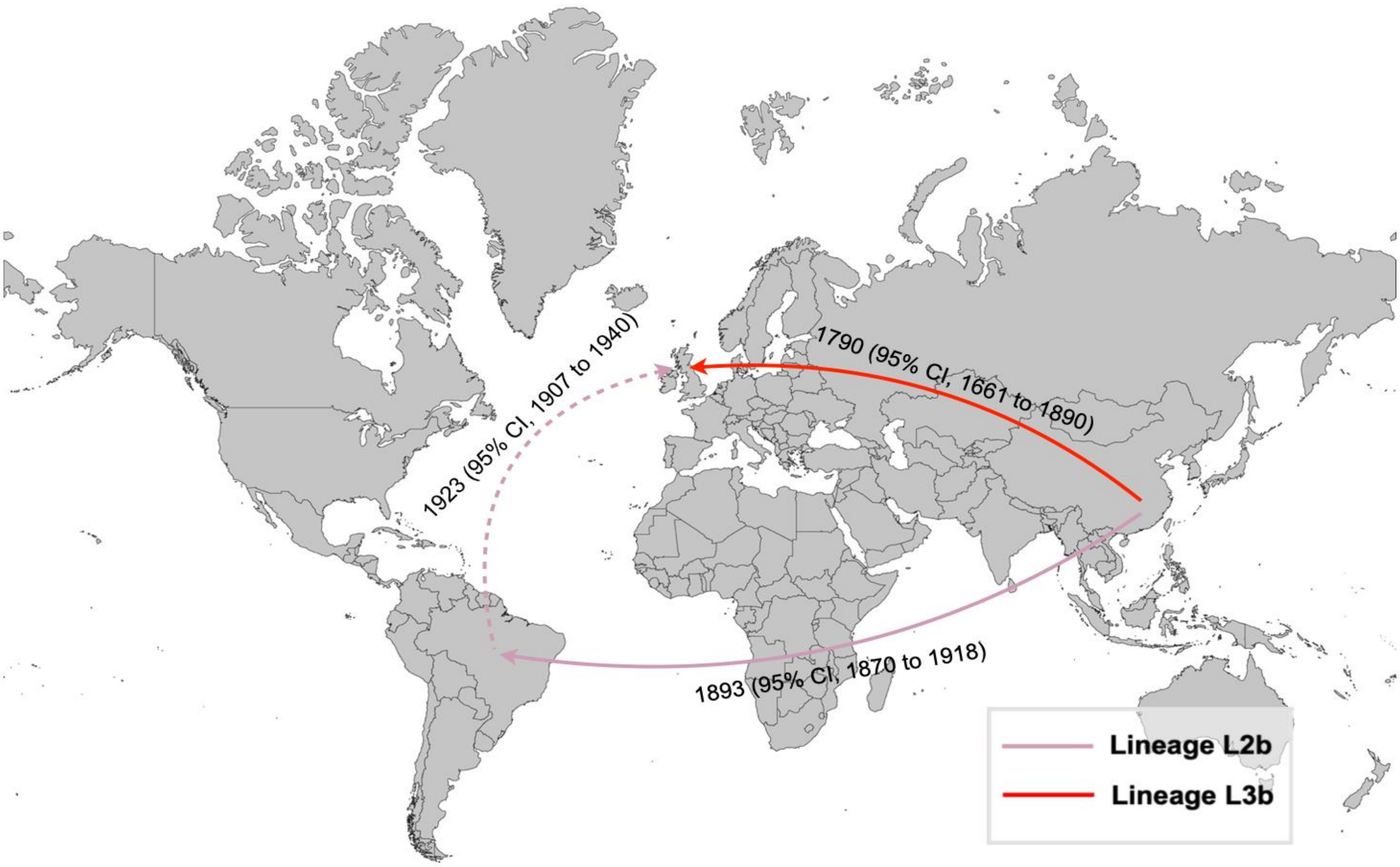


**S7 Fig. Historical international transmissions of bvSP lineages L2b and L3b are depicted with arrows representing the transmission paths. The pink and red lines represent L2b and L3b, respectively.**

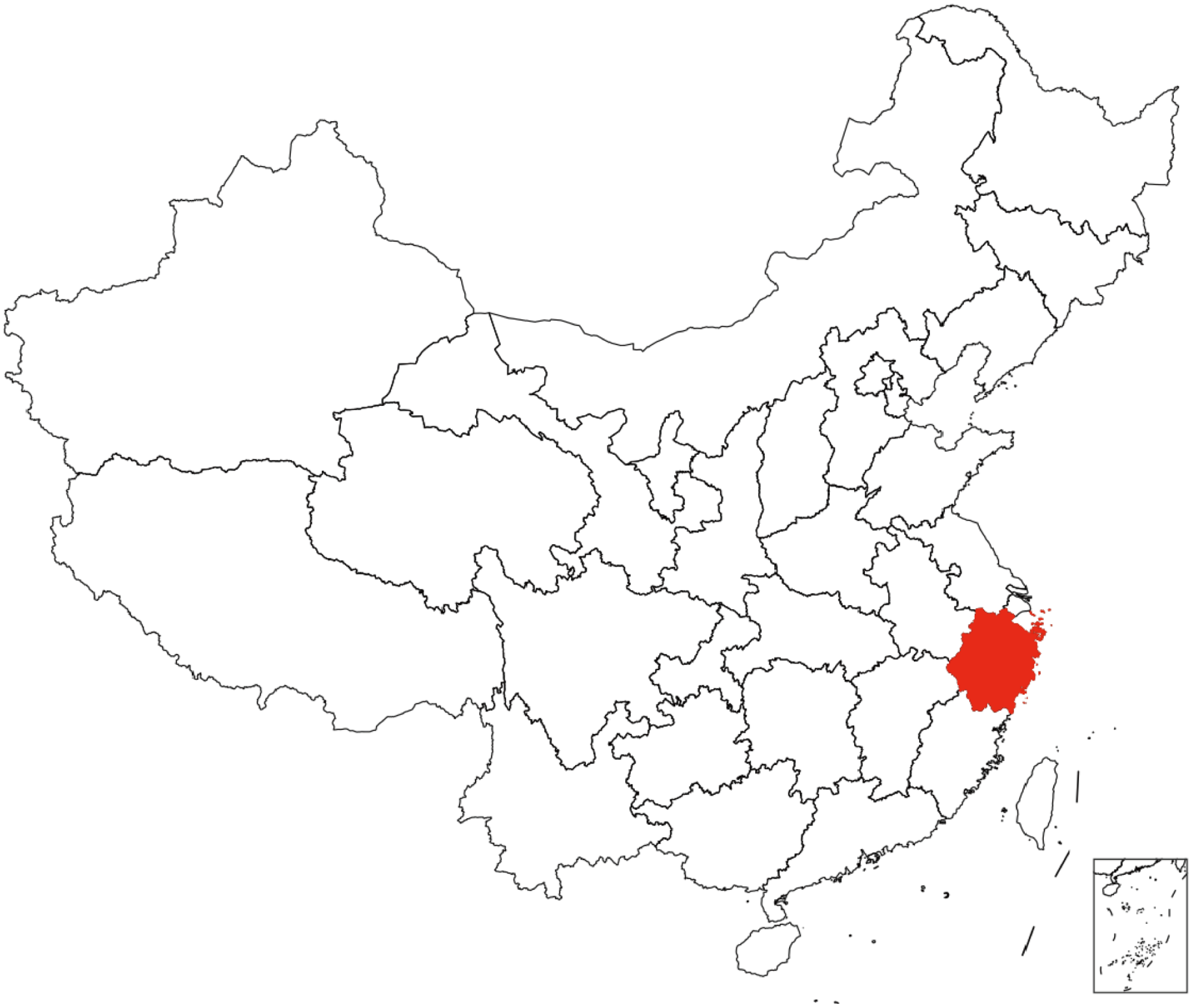

**S8 Fig. Potential transmission events (n=53) of S. Gallinarum biovar Pullorum (bvSP).** The cgSNP distances were calculated between the bvSP strains isolated from Zhejiang Province (n=95) and those from China with available provincial information (n=435). Only cgSNP distances less than two are depicted, with darker colors indicating a higher transmission event.

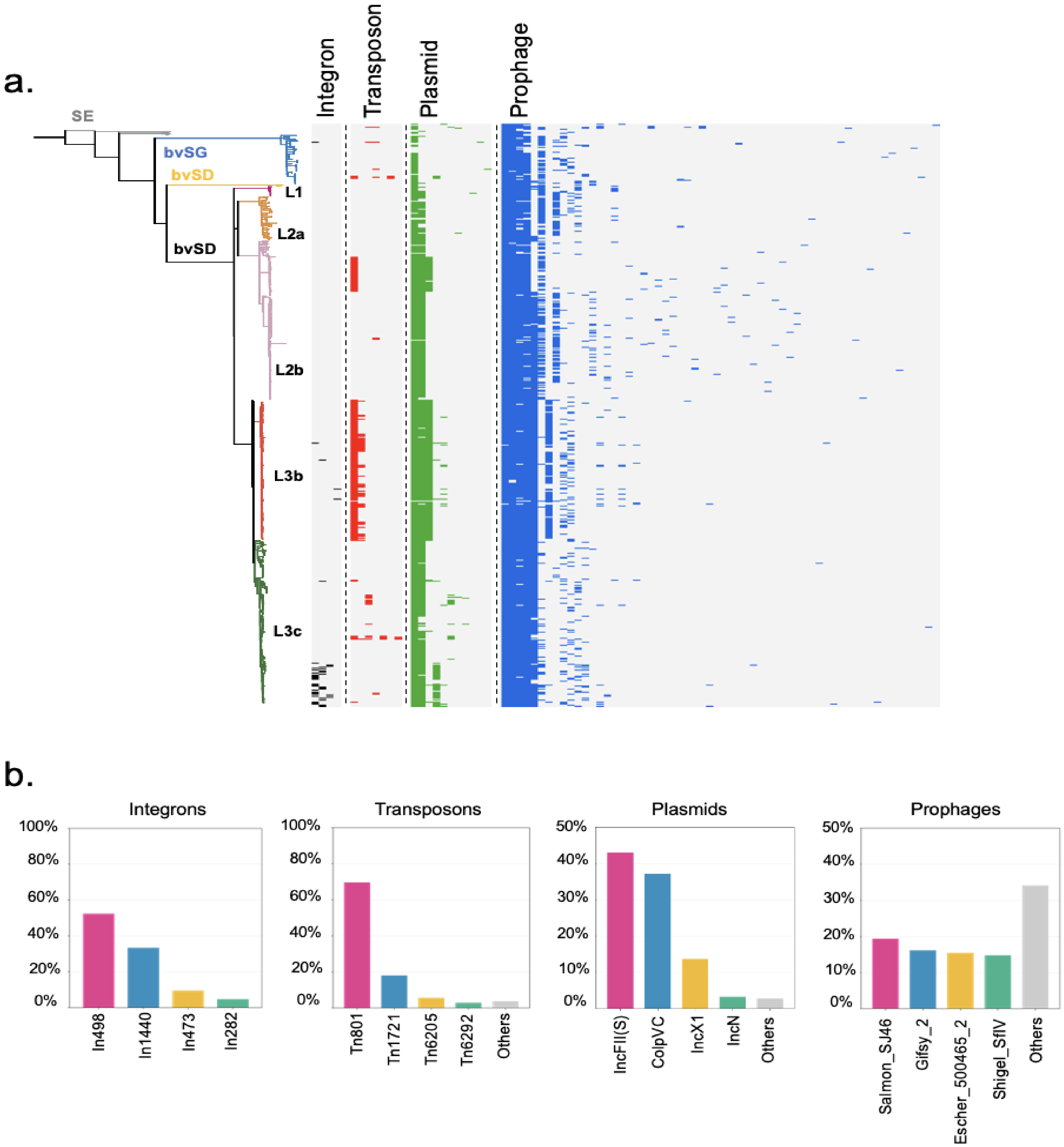


**S9 Fig. The carriage of four predominant mobilome.** (a). The phylogenetic tree of *S.* Gallinarum was constructed using cgSNPs, with distinct colors representing each *S.* Gallinarum biovar; *Salmonella* serovar Enteritidis (SE.) is depicted in gray. Furthermore, different colors are assigned to represent various lineages of bvSP: fuchsia for L1, orange for L2a, pink for L2b, red for L3b, and green for L3c. Heatmaps on the right side illustrate the presence of integrons, transposons, plasmids, and prophages carried by the corresponding *Salmonella* strains. (b). Predominant types of mobilomes prevalent among bvSP are depicted. The x-axis of the bar graph illustrates the top five mobilome types in bvSP based on the total count within each category.

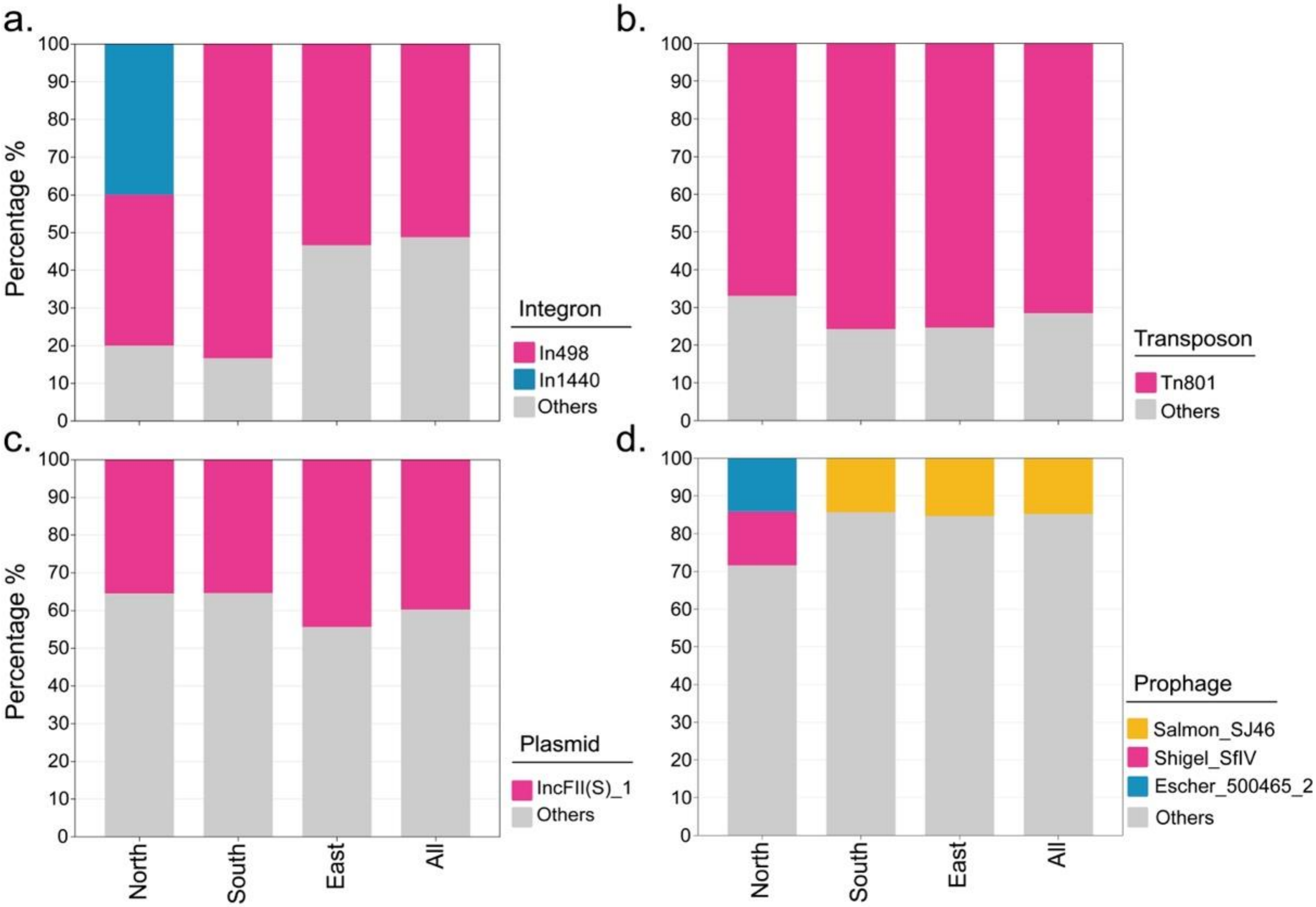


**S10 Fig. Types of predominant mobile genetic elements carried by various regions.** From a-d presents integron, transposon, plasmid, and prophage, respectively.

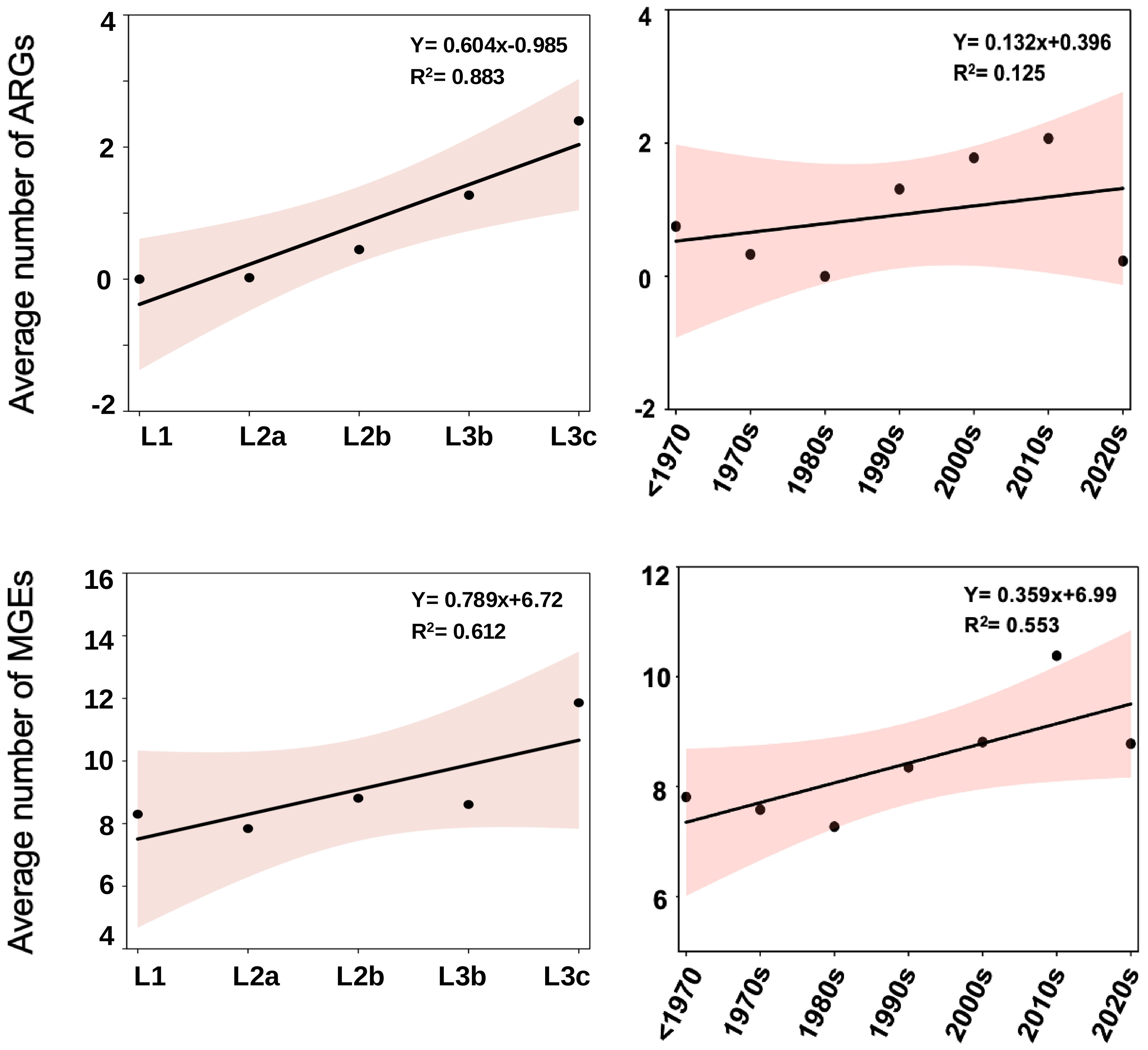


**S11 Fig. Trends in both resistome and mobilome quantities over time and across lineages.**

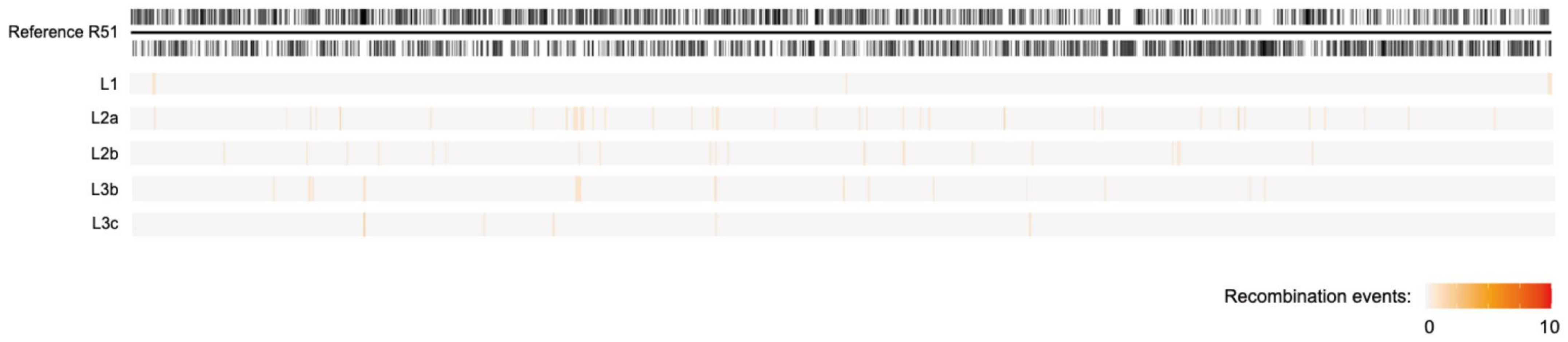


**S12 Fig. Recombination removal using Gubbins.** Recombination in five lineages (L1, L2a, L2b, L3b, L3c) were removed using Gubbins with default parameters. The recombination regions for each lineage were mapped onto the reference genome, *S*. Gallinarum R51. Different colors represent the number of recombination events in each *S*. Gallinarum lineages strains, with darker colors indicating higher frequencies of recombination.
